# Secretome-Mediated Antimicrobial and Immunomodulatory Activity of *Lactobacillus johnsonii* Against Multidrug-Resistant *Enteroaggregative Escherichia coli*

**DOI:** 10.64898/2026.04.02.716048

**Authors:** Sai Madhuri Vasamsetti, Sheeba M, Yasaswi Khaderbad, Anubha G, Vijay Morampudi

**Author notes:** Corresponding author: (VM).

## Abstract

*Enteroaggregative Escherichia coli* (EAEC) is a leading cause of persistent diarrhea in children in low- and middle-income countries, and the emergence of multidrug-resistant (MDR) strains necessitates non-antibiotic therapeutic strategies. This study evaluates *Lactobacillus johnsonii*, previously characterized by our group, as a probiotic candidate against a clinically confirmed MDR EAEC isolate resistant to ampicillin, ciprofloxacin, azithromycin, amoxicillin, and gentamicin. *L. johnsonii* demonstrated robust gastrointestinal resilience, high cell surface hydrophobicity, phenol tolerance, and rapid autoaggregation reaching 80.4 ± 2.3% by 4 hours, collectively supporting mucosal colonization potential. In antimicrobial assays, *L. johnsonii* produced zones of inhibition against MDR EAEC substantially exceeding those of gentamicin, reduced viable biofilm-associated EAEC by over 80%, and displaced pre-adhered EAEC from HCT-116 intestinal epithelial cells in a time-dependent manner. *L. johnsonii* also attenuated MDR EAEC-induced gas production and reduced nitric oxide levels by 67.7% in infected RAW 264.7 macrophages, suggesting immunomodulatory activity. Nutrient competition did not appear to contribute to EAEC suppression under tested conditions, indicating inhibition is predominantly secretome-dependent. Fractionation of the *L. johnsonii* cell-free supernatant by fast protein liquid chromatography yielded five fractions below 75 kDa; fractions S5 and S6 exhibited sustained bactericidal activity at 6 hours. Gram staining confirmed that both fractions reduced viable EAEC cell numbers, with S6 producing a greater reduction than S5, indicating quantitatively distinct bactericidal potencies. These in vitro findings support the potential of *L. johnsonii* as a biotherapeutic candidate for antibiotic-resistant enteric infections. In vivo validation and chemical characterization of active fractions remain important next steps.

## INTRODUCTION

*Enteroaggregative Escherichia coli* (EAEC) is a recognized cause of acute and persistent diarrhea, particularly among children under five years of age in low- and middle-income countries^1, 2^. The World Health Organization estimates approximately 1.7 billion episodes of childhood diarrhea annually, contributing to over 525,000 deaths each year, with EAEC accounting for a disproportionate share of persistent cases lasting more than 14 days^3, 4^. In resource-limited settings across South Asia and sub-Saharan Africa, EAEC has been identified as one of the most prevalent enteric pathogens in both community and hospital surveillance studies, often surpassing other diarrheagenic *E. coli* pathotypes in frequency^2, 5^. Beyond acute illness, recurrent EAEC infection is associated with growth faltering, wasting, and impaired cognitive development in young children^6, 7^.

EAEC exhibits substantial genomic and virulence heterogeneity across clinical isolates^8^. It colonizes the intestinal mucosa through a characteristic “stacked-brick” aggregative adherence pattern mediated by aggregative adherence fimbriae (AAFs), encoded on the virulence plasmid pAA^3^. Following attachment, EAEC stimulates excessive mucus secretion and forms structured biofilms that facilitate persistent colonization and immune evasion^5, 9^. This biofilm-associated lifestyle directly enables persistence within the host, as the extracellular matrix shields bacteria from antimicrobial peptides and supports sustained enterotoxin secretion and mucosal inflammation^3, 10^. Clinically, EAEC infection presents as watery or mucoid diarrhea with nausea, low-grade fever, and abdominal cramping, with recurrent episodes predominantly affecting young children^6, 7^.

The global rise of multidrug-resistant (MDR) EAEC strains has significantly complicated clinical management. Resistance to ampicillin, ciprofloxacin, and azithromycin has been documented across multiple geographic settings^11^, and biofilm-embedded EAEC can tolerate antibiotic concentrations several-fold above the minimum inhibitory concentration of planktonic cells^5, 9, 12^. This convergence of acquired resistance and biofilm-mediated phenotypic tolerance leaves few reliable treatment options and creates a strong rationale for non-antibiotic strategies capable of targeting both planktonic and biofilm-associated EAEC simultaneously.

Probiotics, particularly *Lactobacillus* species, inhibit enteric pathogens through competitive exclusion, production of organic acids and bacteriocins, and modulation of host immune responses^13, 14^. Among these, *Lactobacillus johnsonii* (*L. johnsonii*) has attracted interest for its acid and bile tolerance, strong mucosal adherence, and capacity to produce diverse low-molecular-weight antimicrobial compounds^15, 16^. Genomic analyses have identified enrichment of phosphotransferase system transporters and proteolytic enzymes in *L. johnsonii*, supporting metabolic adaptability within the gastrointestinal environment^15^. The strain also disrupts pre-formed biofilms through exopolysaccharide-mediated interference^17, 18^, and clinical studies in children have demonstrated safe reduction of *Helicobacter pylori* colonization following supplementation^19^. Activity against *Candida albicans* and methicillin-resistant *Staphylococcus aureus* has further extended its documented antimicrobial spectrum^16, 20^.

Despite its well-documented antimicrobial repertoire, activity of *L. johnsonii* against MDR EAEC has not been investigated, and probiotic-based strategies for this pathotype represent a meaningful gap in the literature^21^. The biofilm-forming, toxin-secreting, and antibiotic-resistant characteristics of this pathotype present a far more complex challenge than previously studied organisms^22^. To address this, the present study evaluated *L. johnsonii* against a clinically confirmed MDR EAEC isolate through a comprehensive set of in vitro assays encompassing gastrointestinal stress tolerance, surface adhesion properties, biofilm inhibition, competitive exclusion and displacement from intestinal epithelial cells, and direct antimicrobial activity. We also employed fast protein liquid chromatography (FPLC) to fractionate the *L. johnsonii* cell-free supernatant and identify bioactive components with sustained bactericidal activity, representing to our knowledge the first FPLC-based secretome characterization of a probiotic against MDR EAEC. Additionally, the capacity of *L. johnsonii* to attenuate EAEC-induced gas production and macrophage-derived nitric oxide was examined to characterize its immunomodulatory potential.

## MATERIALS AND METHODS

### Bacterial strains, media, and growth conditions

*L. johnsonii* was isolated from homemade Indian curd collected in Hyderabad, India. Strain identity was confirmed by PCR amplification using *L. johnsonii*-specific primers, yielding a distinct 126 bp amplicon, followed by 16S rRNA gene sequencing using universal primers (27F: 5′-AGAGTTTGATCCTGGCTCAG-3′ and 1492R: 5′-GGTTACCTTGTTACGACTT-3′). The sequence exhibited greater than 98% similarity to *L. johnsonii* strain 2317 and has been deposited in GenBank under accession number PV739486. This is the same isolate previously characterized and reported by our group in a related study investigating activity against attaching and effacing pathogens^23^. *L. johnsonii* was routinely cultured in de Man, Rogosa, and Sharpe (MRS) broth (Millipore Sigma, Cat. No. 69966) supplemented with 0.05% (w/v) cysteine (Sigma-Aldrich, Cat. No. 168149) at 37°C for 18 to 24 hours. Cysteine supports anaerobic growth and metabolic activity of lactic acid bacteria^24, 25^. The MDR EAEC clinical isolate was obtained from the Asian Institute of Gastroenterology (AIG Hospitals), Hyderabad, India, under a formal Material Transfer Agreement. The MDR phenotype was confirmed at AIG Hospitals using clinical diagnostic kits prior to transfer and independently verified in our laboratory by antibiotic susceptibility profiling as described below. MDR EAEC was cultured in Luria-Bertani (LB) broth (HiMedia, Cat. No. M1245) at 37°C with shaking at 200 rpm. HCT-116 human colorectal epithelial cells (NCCS) were maintained in McCoy’s 5A medium (Gibco, Cat. No. 16600082) supplemented with 10% fetal bovine serum (FBS; Gibco, Cat. No. 10270-106) at 37°C under 5% CO_2_. HCT-116 is a non-mucus-producing human intestinal epithelial cell line, making it suitable for direct bacteria-epithelial interaction studies without interference from mucus layers. RAW 264.7 murine macrophage cells were cultured in Dulbecco’s Modified Eagle Medium (DMEM) supplemented with 10% FBS and 1X penicillin-streptomycin at 37°C under 5% CO_2_.

### Growth curve analysis and colony-forming unit determination

*L. johnsonii* was cultured in 5 mL of MRS supplemented with 0.05% (w/v) cysteine (MRS+), while MDR EAEC was cultured in 5 mL of LB broth, both at 37°C with shaking at 200 rpm. Optical density at 600 nm (OD_600_) was recorded at 0, 2, 4, 8, 24, and 48 hours using a BioSpectrometer Basic (Molecular Devices). For colony-forming unit (CFU) determination, overnight cultures were adjusted to OD_600_ = 1.0 and serially diluted tenfold from 10^-1^ to 10^-7^. From each dilution, 10 µL was plated in triplicate on MRS+ agar for *L. johnsonii* and LB agar for MDR EAEC, and plates were incubated overnight at 37°C. Viable counts were calculated as: CFU/mL = (Number of colonies × dilution factor) / volume plated (mL).

### Antibiotic susceptibility profiling of MDR EAEC

The antibiotic resistance profile of the EAEC isolate was determined by broth microdilution. MDR EAEC was standardized to OD_600_ = 1.0 and exposed to six antibiotics: ampicillin, ciprofloxacin, azithromycin, amoxicillin, gentamicin, and cefotaxime, each at concentrations of 6.25, 12.5, 25, 50, and 100 µg/mL in LB broth in a 96-well plate format. Uninoculated LB broth served as the sterility control and untreated MDR EAEC as the growth control. Plates were incubated at 37°C for 24 hours, after which OD_600_ was measured. Isolates showing no meaningful growth inhibition (less than 50% reduction in OD600 relative to untreated control) to agents from three or more antimicrobial classes were classified as phenotypically MDR in accordance with the categorical framework of Magiorakos et al.^26^. It should be noted that formal MIC determination was not performed and antibiotic classification was based on OD600-based growth inhibition as a surrogate measure.

### Assessment of gastrointestinal stress tolerance

Acid tolerance: *L. johnsonii* was inoculated into MRS+ broth adjusted to pH 1.2, 1.5, 2.0, and 2.5 using 1 M hydrochloric acid (HiMedia) and incubated at 37°C for up to 3 hours. Samples were collected at 0, 1, 2, and 3 hours and viability assessed by measuring OD600 using a BioSpectrometer Basic (Molecular Devices). Uninoculated MRS+ broth at each pH served as the sterility control; *L. johnsonii* at pH 6.5 served as the positive growth control. Bile salt tolerance: *L. johnsonii* was inoculated into MRS+ broth supplemented with 0.3% (w/v) bile salts (HiMedia, Cat. No. CR010) and incubated at 37°C for 3 hours. Growth was monitored by measuring OD600 at 0, 1, 2, and 3 hours. Simulated gastric and intestinal fluid tolerance was assessed as previously described^23^.

### Surface adhesion properties: Hydrophobicity, Phenol Tolerance, and Autoaggregation

Cell surface hydrophobicity: The microbial adhesion to hydrocarbons (MATH) assay was performed as described by Rosenberg et al.^27^. Briefly, overnight *L. johnsonii* cultures were resuspended in PBS to OD_600_ = 1.0. One milliliter of xylene or chloroform (SRL) was added to 3 mL of the suspension, vortexed for 2 minutes, and incubated at 37°C for 15 minutes for phase separation. The aqueous phase absorbance was measured at 600 nm and hydrophobicity calculated as: Hydrophobicity (%) = [1 - (A_final_ / A_initial_)] × 100. Phenol tolerance: *L. johnsonii* (10^7^ CFU/mL) was inoculated into MRS+ broth containing 0.2% or 0.4% (w/v) phenol and incubated at 37°C for 6 hours. Viable counts were determined by plating on MRS+ agar and expressed as CFU/mL. Phenol tolerance was assessed as a proxy for resistance to aromatic fermentation metabolites that accumulate in the colon under dysbiotic conditions^28^. Autoaggregation: Overnight *L. johnsonii* culture was washed, resuspended in PBS to OD_600_ = 0.5 ± 0.05, and incubated at room temperature without agitation. Aliquots (100 µL) of the upper phase were collected at 0, 4, and 24 hours, and autoaggregation calculated as: Autoaggregation (%) = [1 - (A_t_ / A_0_)] × 100.

### Agar overlay assay

*L. johnsonii* was spot-inoculated onto MRS+ agar plates (10 µL, approximately 10^5^ CFU/spot) and incubated at 37°C for 48 hours. Plates were then overlaid with LB agar containing 10^7^ CFU of MDR EAEC per plate. After solidification, plates were incubated at 37°C for 24 hours. The zone of inhibition (ZOI) was measured in centimeters and compared to gentamicin (30 µg/mL) as the antibiotic control. MDR EAEC overlaid on MRS+ agar without *L. johnsonii* served as the negative control. All assays were performed in triplicate and repeated independently three times.

### Biofilm inhibition assay

MDR EAEC and *L. johnsonii* were inoculated individually and in combination into 96-well plates containing 180 µL of Brain Heart Infusion (BHI) broth (20 µL inoculum per well). Heat-killed *L. johnsonii* (HK-LJ), prepared by incubation at 80°C for 30 minutes with complete killing confirmed by plating, was included as a viability-dependent control. Each condition was tested in six replicate wells. After 48 hours at 37°C, wells were washed, stained with 0.1% crystal violet for 30 minutes, washed again, air dried, and solubilized with 100 µL of 100% ethanol. Absorbance was measured at 570 nm (Molecular Devices). Viable bacteria within biofilms were quantified by scraping, resuspending in PBS, serially diluting, and plating on MRS+ agar for *L. johnsonii* and LB agar for MDR EAEC. Uninoculated BHI broth served as the sterility control.

### Adherence and protection assays

HCT-116 cells were seeded in 24-well plates at 2.5 × 10^4^ cells per well and grown to 90% confluency. For the adherence assay, monolayers were incubated with *L. johnsonii* or MDR EAEC at MOI 1:25 for 3 hours at 37°C under 5% CO_2_. After washing three times with PBS, monolayers were lysed with 1% Triton X-100 for 10 minutes, and serial dilutions plated on the appropriate agar media. Adherence was expressed as a percentage of the inoculum recovered. Three protection assays were conducted at MOI 1:25: Exclusion: monolayers pre-incubated with *L. johnsonii* for 3 hours, then challenged with MDR EAEC for 3 hours; Displacement: monolayers infected with MDR EAEC for 3 hours, then treated with *L. johnsonii* for 3 hours; Competition: *L. johnsonii* and MDR EAEC co-inoculated simultaneously for 6 hours. After each assay, wells were washed and lysed, and CFU enumerated. *L. johnsonii* growth was monitored throughout by plating on MRS+ agar.

### Gas production assay

Overnight cultures of MDR EAEC and *L. johnsonii* were adjusted to OD_600_ = 1.0. MDR EAEC (1 × 10^8^ CFU in 20 µL) was inoculated into the lower third of glass tubes containing 5 mL of 0.7% LB soft agar. Three milliliters of 0.7% MRS+ soft agar containing *L. johnsonii* (1 × 10^8^ CFU) was then layered on top. Controls included: (i) MDR EAEC in LB soft agar overlaid with MRS+ soft agar without *L. johnsonii* to confirm EAEC gas production; and (ii) *L. johnsonii* in MRS+ soft agar overlaid with plain LB soft agar to confirm absence of probiotic gas production. All tubes were incubated at 37°C for 24 hours under anaerobic conditions. Gas production was recorded qualitatively as visible agar splitting or displacement.

### Nitric oxide detection in MDR EAEC-infected macrophages

The effect of *L. johnsonii* on nitric oxide (NO) production was assessed using the Griess reagent assay. RAW 264.7 cells were seeded at 1 × 10^4^ cells per well in 96-well plates and allowed to adhere overnight. Cells were infected with MDR EAEC at MOI 1:10 or stimulated with LPS (500 ng/mL) for 24 hours, followed by addition of *L. johnsonii* at MOI 1:25 for 6 hours. Supernatants were collected and mixed with Griess reagent (1% sulfanilamide and 0.1% naphthylethylenediamine dihydrochloride in 5% phosphoric acid) for 10 minutes at room temperature. Absorbance was measured at 540 nm, and nitrite concentrations calculated from a sodium nitrite standard curve and expressed as µM nitrite per well. Uninfected untreated cells served as the basal control.

### Nutrient competition assay

Overnight cultures of both strains were adjusted to 10^7^ CFU/mL. *L. johnsonii* and MDR EAEC were co-inoculated simultaneously into 24-well plates at 10^7^ CFU per organism per well in either McCoy’s 5A incomplete medium supplemented with 2.5% FBS (nutrient-rich) or sterile PBS (nutrient-limited), and incubated at 37°C for 6 hours. Single-organism cultures served as controls. After incubation, 10 µL of serially diluted samples were plated on the appropriate agar media. CFU/mL were determined after overnight incubation at 37°C. The assay was performed in triplicate and repeated independently three times.

### Preparation of bacterial cell-free supernatant and lysate

Bacterial cell-free supernatant (CFS) was prepared by culturing *L. johnsonii* in MRS+ broth for 18 to 24 hours, centrifuging at 3,500 rpm for 20 minutes, and filtering the supernatant through a 0.22 µm membrane (Sartorius). The CFS was used without pH neutralization; the native pH of the CFS was approximately 4.2, which may contribute to the observed antimicrobial activity. For lysate preparation, 10 mL of overnight culture was pelleted, resuspended in lysozyme solution (10 mg/mL in Tris-EDTA buffer, pH 8.0), and incubated at 37°C for 2 hours. Cells were sonicated using a Sonopulse Probe (25 cycles of 15 seconds each with rest intervals on ice) and filtered through a 0.22 µm membrane (Millipore). Protein concentration in both preparations was quantified using the DC Protein Assay (Bio-Rad) with BSA as the standard. MDR EAEC was incubated with 12.5, 25, or 50 µg protein per 100 µL of lysate or CFS for 6 hours at 37°C, and viable counts were determined by CFU enumeration.

### Fast protein liquid chromatography

*L. johnsonii* was cultured in 1 L of MRS+ broth at 37°C with shaking at 200 rpm for 48 hours. The culture was centrifuged at 3,500 rpm (Hitachi) and the supernatant filtered through a 0.22 µm cellulose nitrate membrane (Sartorius). The clarified supernatant was maintained at 8°C with continuous stirring overnight. Ammonium sulfate (SRL, Cat. No. 82126) was added to 30% (w/v) final concentration and the solution incubated at 8°C for 18 hours. The precipitate was collected by centrifugation at 12,000 × g for 15 minutes at 6°C, resuspended in 50 mM phosphate buffer (pH 7.0), and concentrated by lyophilization (Scanvac) overnight. Approximately 5 mg of protein in 500 µL of phosphate buffer was subjected to FPLC using a Superdex 75 size-exclusion column (Cytiva) at a flow rate of 0.5 mL/min. Five fractions (S1, S2, S4, S5, and S6) were collected; fraction S3 was not collected as it eluted within a poorly resolved region of the chromatogram with insufficient UV signal for reliable fraction definition. Each fraction was lyophilized and reconstituted in sterile PBS at 30 µg/mL for antimicrobial activity testing against MDR EAEC at 1 and 6 hours. Fractions showing significant inhibition at both time points were selected for further characterization by Gram staining.

### Gram staining

Gram staining was performed to morphologically confirm and visually quantify the bactericidal activity of L. johnsonii CFS (25 and 50 µg/mL) and fractions S5 and S6 (30 µg/mL each) against MDR EAEC following 6 hours of treatment at 37°C. Untreated MDR EAEC served as the control. Thin smears were prepared, air dried, heat fixed, and stained following standard Gram protocol. Preparations were examined under oil immersion at 100× magnification, and changes in cell density, morphology, and staining intensity were recorded.

### Statistical analysis

All experiments were performed with a minimum of three independent biological replicates, each conducted in technical triplicate, unless otherwise stated. Data are presented as mean ± standard error of the mean (SEM). Statistical analyses were performed using GraphPad Prism version 8.4.2 (GraphPad Software, San Diego, USA). For comparisons between two groups, an unpaired two-tailed Student’s t-test was used. For comparisons among three or more groups, one-way ANOVA with Dunnett’s multiple comparisons test was applied. For experiments with two independent variables, two-way ANOVA with Dunnett’s multiple comparisons test was applied. A p-value less than 0.05 was considered statistically significant. Significance thresholds are denoted in figures as follows: p < 0.05 (*), p < 0.01 (**), p < 0.001 (***), and p < 0.0001 (****); ns indicates no statistical significance.

## RESULTS

### Confirmation of MDR phenotype in the EAEC isolate

The EAEC isolate was obtained from a clinical source, and its multidrug-resistant (MDR) status was independently verified prior to experimentation. The isolate exhibited robust growth in LB broth, reaching stationary phase by 24 hours (Figure 1A). Antibiotic susceptibility profiling showed no meaningful growth inhibition in response to five first-line agents (ampicillin, ciprofloxacin, azithromycin, amoxicillin, and gentamicin) across the tested concentration range (6.25 to 100 µg/mL). Cefotaxime, included as a sixth comparator, was the only agent to produce concentration-dependent inhibition (Figure 1B). The lack of appreciable inhibition across agents representing four distinct antibiotic classes (penicillins, fluoroquinolones, macrolides, and aminoglycosides) is consistent with an MDR phenotype as defined by Magiorakos et al.^26^, noting that the MDR status of the isolate was also established by the supplying institution using standard clinical diagnostic methods. These observations support the use of this isolate for evaluating non-antibiotic therapeutic strategies.

**Figure 1.**
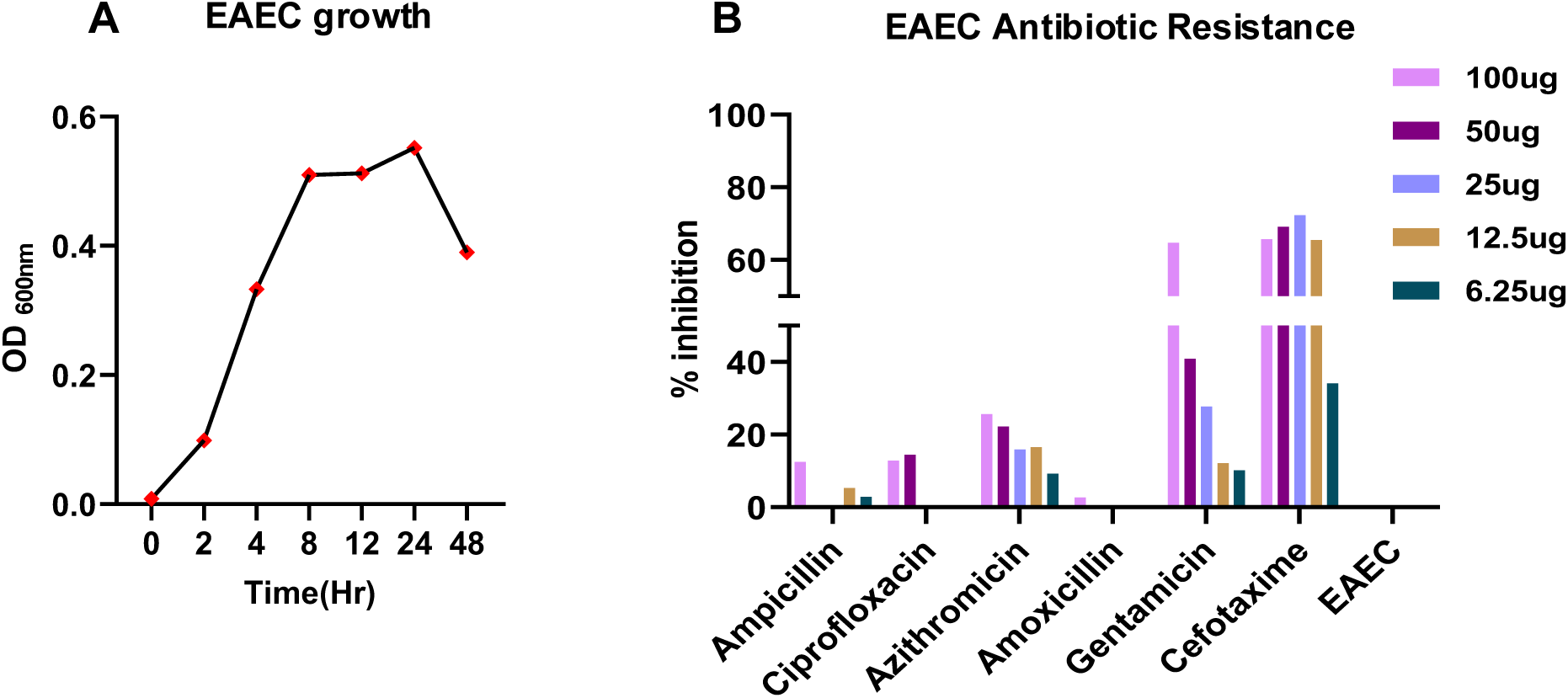
Confirmation of the MDR phenotype of the EAEC clinical isolate. (A) Growth kinetics of MDR EAEC in LB broth measured by OD600 at 0, 2, 4, 8, 24, and 48 hours. (B) Antibiotic susceptibility profile of MDR EAEC following 24 hours of exposure to ampicillin, ciprofloxacin, azithromycin, amoxicillin, gentamicin, and cefotaxime at 6.25, 12.5, 25, 50, and 100 µg/mL. Untreated MDR EAEC served as the growth control and uninoculated LB broth as the sterility control. Results expressed as OD600. Data are mean ± SEM (n = 3 independent biological replicates). Statistical analysis: one-way ANOVA with Dunnett’s multiple comparisons test. Significance: p < 0.05 (*), p < 0.01 (**), p < 0.001 (***), p < 0.0001 (****); ns, not significant.

### Probiotic characterization of *L. johnsonii*

To evaluate the suitability of *L. johnsonii* as a probiotic candidate against MDR EAEC, its gastrointestinal stress tolerance and surface adhesion properties were assessed. *L. johnsonii* grew consistently in MRS+ broth over 48 hours, with a well-defined exponential phase between 2 and 8 hours followed by entry into stationary phase (Figure 2A). The strain demonstrated robust acid tolerance, maintaining viability at pH 1.2 to 2.0 for up to 1 hour and sustaining growth at pH 2.5 for the full 3-hour incubation period (Figure 2B). Bile salt tolerance at 0.3% and resistance to simulated gastric and intestinal fluids containing pepsin and trypsin have been characterized for this isolate using CFU-based viability assays and are reported in detail elsewhere^23^. OD600-based confirmation of bile salt tolerance is shown in Figure 2C. *L. johnsonii* retained significant viability at both 0.2% and 0.4% phenol relative to the untreated MRS+ control, with a significant reduction at 0.4% (Figure 2D, p < 0.001). Cell surface hydrophobicity was 74.7 ± 3.6% with xylene and 84.4 ± 5.5% with chloroform (Figure 2E), consistent with strong epithelial adhesion potential. Autoaggregation increased rapidly to 80.4 ± 2.3% by 4 hours and remained stable at 24 hours (Figure 2F, p < 0.0001 versus 0 hours). Collectively, these findings establish that *L. johnsonii* possesses the gastrointestinal resilience and surface adhesion properties required for effective mucosal colonization and persistence.

**Figure 2.**
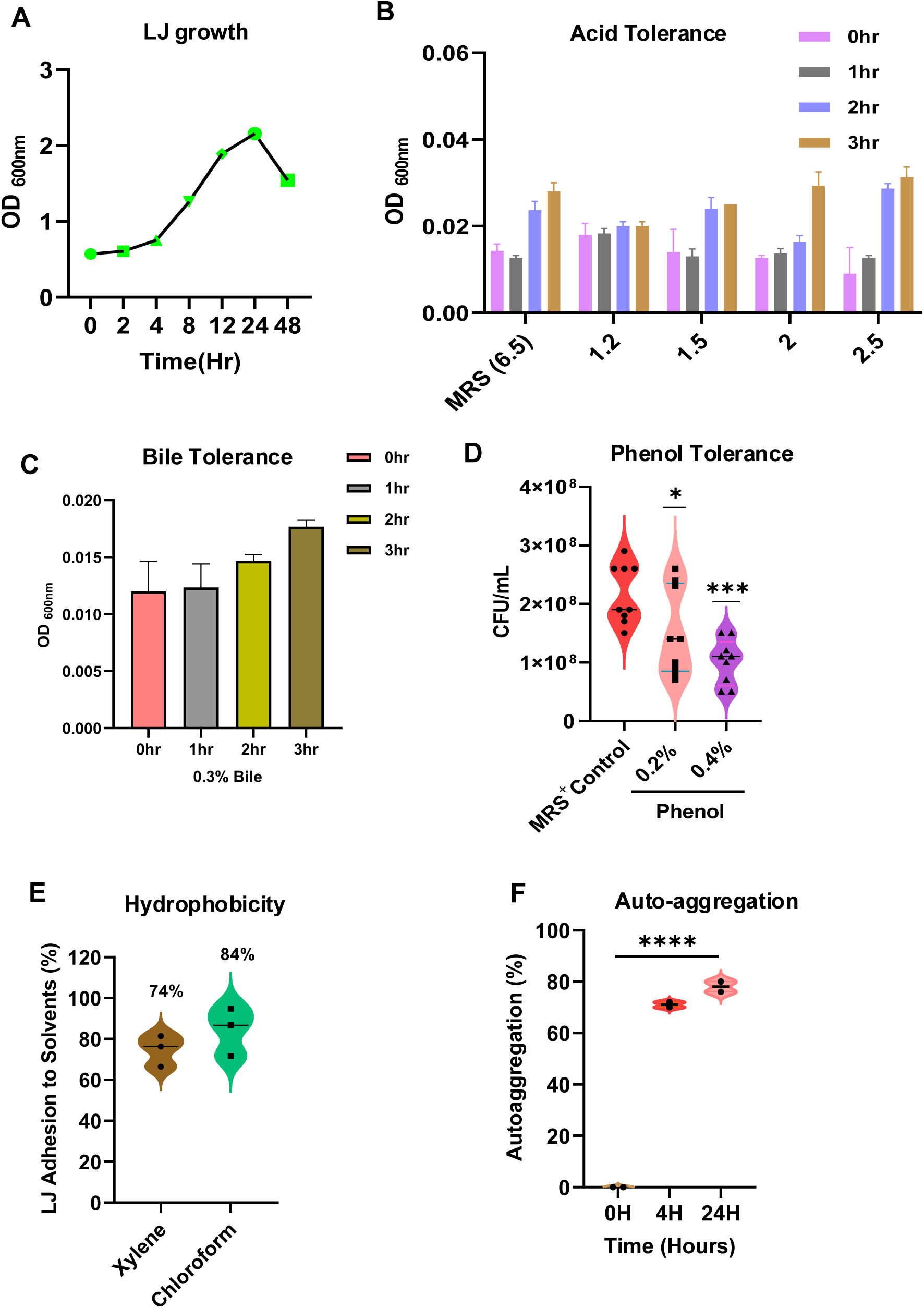
Probiotic characterization of *L. johnsonii*: gastrointestinal stress tolerance and surface adhesion properties. (A) Growth kinetics of *L. johnsonii* in MRS+ broth (OD600) at 0, 2, 4, 8, 24, and 48 hours. (B) Acid tolerance at pH 1.2, 1.5, 2.0, and 2.5 over 0 to 3 hours. Growth at pH 6.5 served as the positive control. Results expressed as OD600. (C) Bile salt tolerance at 0.3% (w/v) bile salts over 0 to 3 hours at 37°C. Results expressed as OD600. CFU-based characterization of bile salt tolerance and simulated gastric and intestinal fluid resistance have been previously reported^23^. (D) Phenol tolerance at 0.2% and 0.4% (w/v) for 6 hours at 37°C. *L. johnsonii* in phenol-free MRS+ broth served as the control. Results expressed as CFU/mL. (E) Cell surface hydrophobicity by MATH assay using xylene and chloroform. Results expressed as percentage adhesion. (F) Autoaggregation measured at 0, 4, and 24 hours. Results expressed as percentage autoaggregation relative to absorbance at time zero. All experiments performed in triplicate with three independent biological replicates (n = 3). Data are mean ± SEM. Statistical analysis: one-way ANOVA with Dunnett’s multiple comparisons test for panel D and unpaired t-test for panels E and F. Significance: p < 0.05 (*), p < 0.01 (**), p < 0.001 (***), p < 0.0001 (****); ns, not significant.

### *L. johnsonii* inhibits MDR EAEC growth and disrupts biofilm formation

The antimicrobial and anti-biofilm activity of *L. johnsonii* against MDR EAEC was evaluated by agar overlay and crystal violet biofilm assays. In the agar overlay assay, *L. johnsonii* produced a zone of inhibition of 3.37 ± 0.17 cm, substantially exceeding that of gentamicin at 30 µg/mL (0.37 ± 0.05 cm), while no inhibition was observed with EAEC alone (Figures 3A and 3B), demonstrating potent contact-independent antimicrobial activity against an isolate to which gentamicin offered no clinical utility. To assess anti-biofilm activity, MDR EAEC was co-cultured with live or heat-killed *L. johnsonii* (HK-LJ) for 48 hours. Crystal violet staining revealed that live *L. johnsonii* significantly reduced EAEC biofilm biomass compared to EAEC alone, while heat-killed *L. johnsonii* (HK-LJ) produced no significant reduction, indicating that biofilm inhibition is dependent on bacterial viability and active metabolic secretion (Figure 3C). CFU enumeration within biofilms confirmed an 81.4% reduction in recoverable EAEC in the presence of live *L. johnsonii*, whereas HK-LJ did not significantly alter viable EAEC counts relative to the untreated biofilm control (Figure 3D), together demonstrating that structural cell components alone are insufficient for anti-biofilm efficacy. Notably, co-culture with MDR EAEC reduced *L. johnsonii* biofilm formation by 71.3% relative to probiotic cultured alone, suggesting reciprocal competitive pressure in the biofilm niche with potential implications for the timing of probiotic administration (Figure 3E).

**Figure 3.**
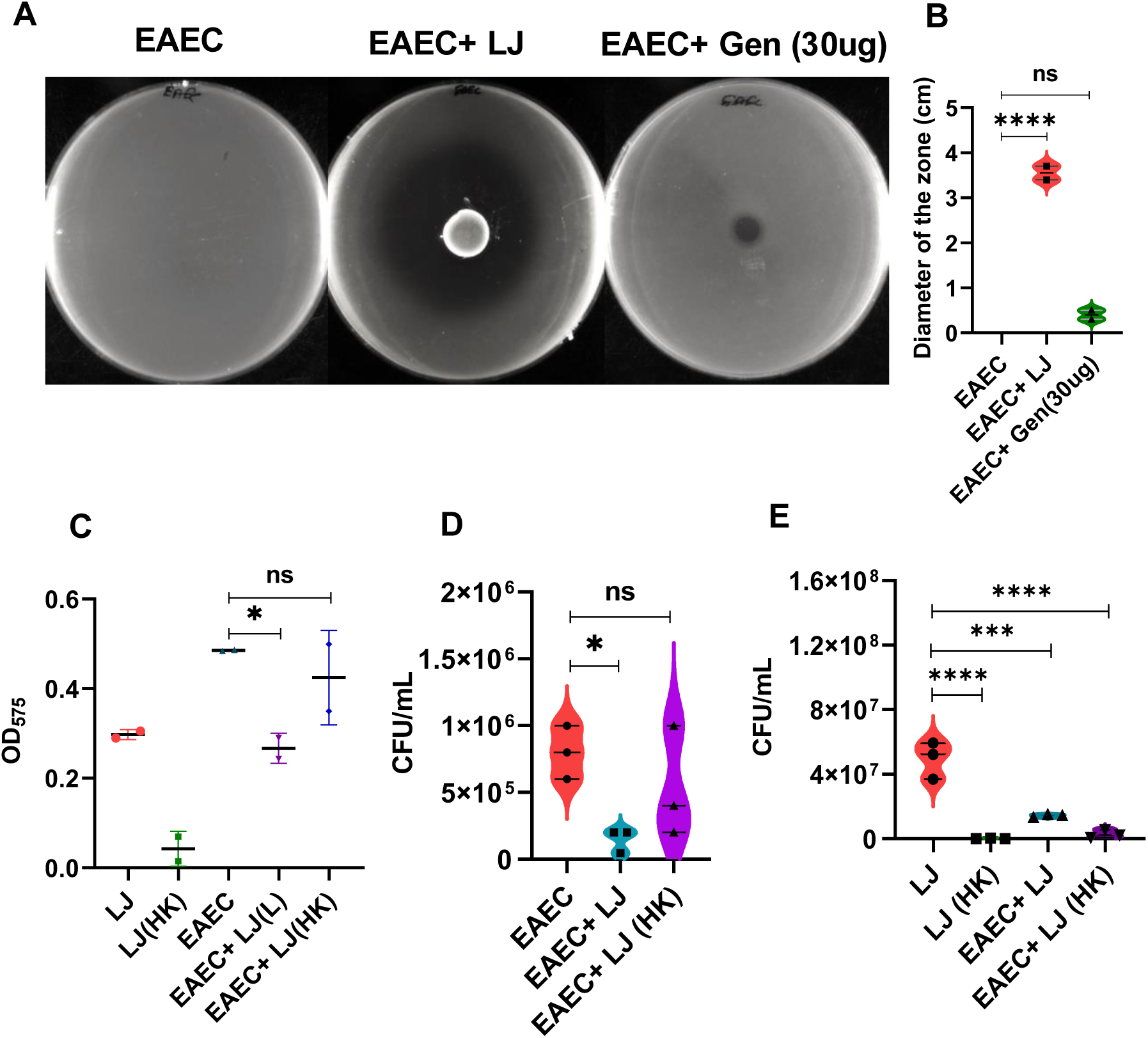
*L. johnsonii* inhibits MDR EAEC growth and disrupts biofilm formation. (A) Representative agar overlay plates showing zones of inhibition for MDR EAEC alone, MDR EAEC overlaid on *L. johnsonii* (LJ), and MDR EAEC treated with gentamicin (30 µg/mL). (B) Quantification of zone of inhibition diameters (cm). (C) Crystal violet staining quantifying MDR EAEC biofilm biomass (OD570) after 48 hours co-culture with live LJ, heat-killed LJ (HK-LJ; prepared at 80°C for 30 minutes, killing confirmed by plating), or no probiotic. (D) CFU enumeration of viable MDR EAEC recovered from biofilms after 48 hours co-culture with LJ or HK-LJ. (E) CFU enumeration of viable L. johnsonii recovered from biofilms after 48 hours co-culture with MDR EAEC. All experiments performed with six technical replicates per condition and three independent biological replicates (n = 3). Data are mean ± SEM. Statistical analysis: one-way ANOVA with Dunnett’s multiple comparisons test. Significance: p < 0.05 (*), p < 0.01 (**), p < 0.001 (***), p < 0.0001 (****); ns, not significant.

### *L. johnsonii* adheres preferentially to intestinal epithelial cells and displaces MDR EAEC through exclusion and displacement

The ability of *L. johnsonii* to adhere to HCT-116 intestinal epithelial cells and competitively inhibit MDR EAEC colonization was assessed through adherence and protection assays at MOI 1:25. *L. johnsonii* adhered to HCT-116 cells at a rate of 3.35% of the inoculum^23^, nearly double the adherence rate of EAEC (1.9%; Figure 4A), indicating a competitive advantage for epithelial niche occupancy. Across the three protection assays, a clear hierarchy of inhibitory efficacy emerged. Pre-incubation of epithelial cells with *L. johnsonii* for 3 hours prior to EAEC challenge significantly reduced EAEC colonization by 59.6% (p < 0.05; Figure 4B), demonstrating that prior epithelial establishment effectively impedes initial EAEC attachment. Post-infection addition of *L. johnsonii* to pre-established EAEC infection displaced adherent pathogens by 34.7% (p < 0.05; Figure 4C), indicating active displacement from the epithelial surface. In contrast, simultaneous co-inoculation produced no significant reduction in EAEC counts (Figure 4D), consistent with a time-dependent colonization resistance mechanism that requires prior epithelial occupancy. Monitoring of *L. johnsonii* growth during each assay revealed significantly enhanced probiotic proliferation under the exclusion condition relative to *L. johnsonii* alone (p < 0.05; Figure 4E), suggesting a growth advantage conferred by prior surface establishment. No significant differences in *L. johnsonii* counts were observed under displacement or competition conditions (Figures 4F and 4G), indicating that EAEC co-culture does not compromise probiotic viability.

**Figure 4.**
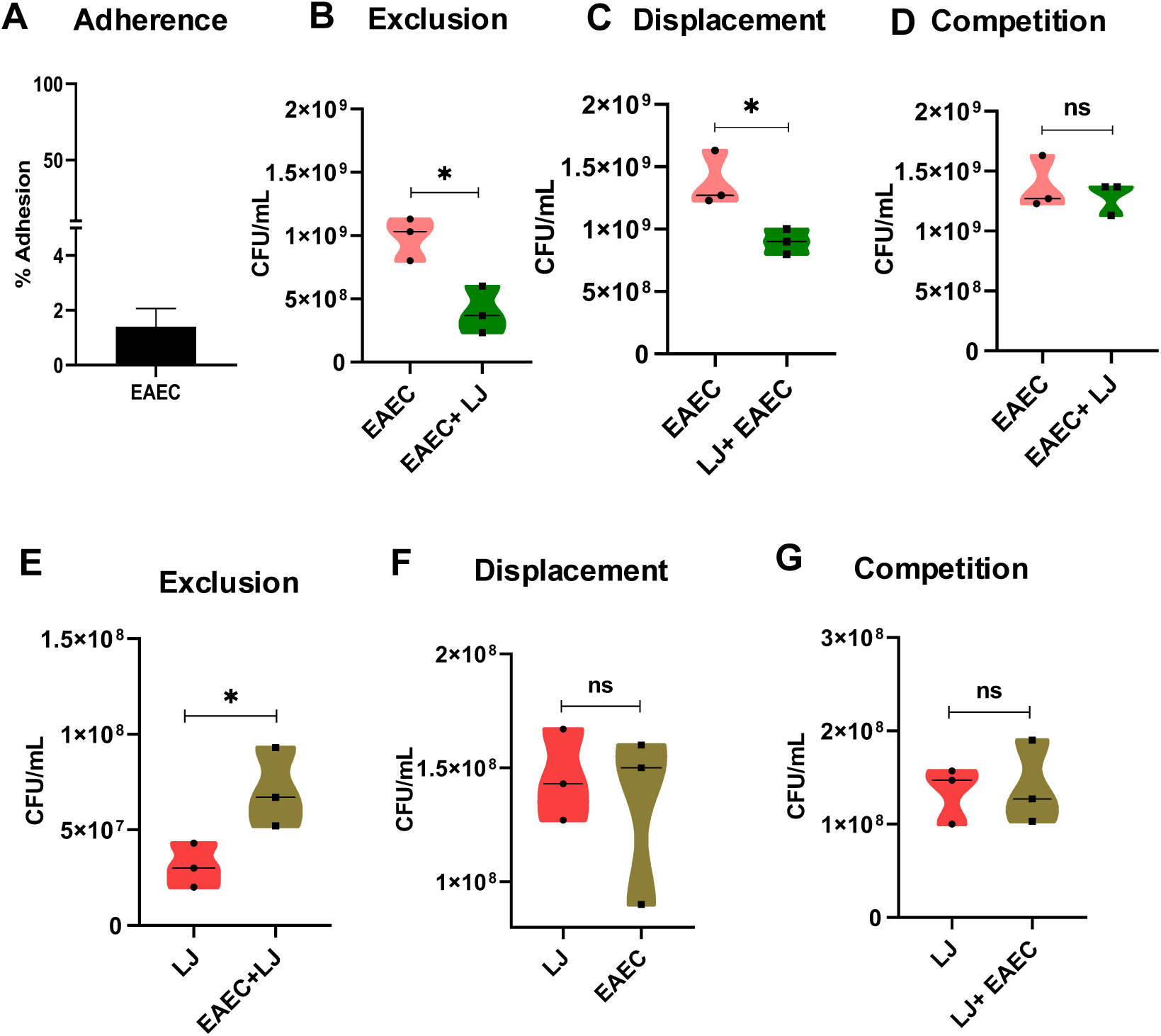
*L. johnsonii* adheres preferentially to intestinal epithelial cells and reduces MDR EAEC colonization through exclusion and displacement. (A) Adherence of MDR EAEC to HCT-116 cells after 3 hours at MOI 1:25, expressed as percentage of inoculum recovered by CFU enumeration after lysis with 1% Triton X-100. (B) Exclusion assay: HCT-116 monolayers pre-incubated with LJ for 3 hours, then challenged with MDR EAEC for 3 hours. MDR EAEC CFU/mL recovered from the adherent fraction. (C) Displacement assay: HCT-116 monolayers infected with MDR EAEC for 3 hours, then treated with LJ for 3 hours. MDR EAEC CFU/mL recovered. (D) Competition assay: LJ and MDR EAEC co-inoculated simultaneously for 6 hours. MDR EAEC CFU/mL recovered. (E–G) Growth of LJ during exclusion (E), displacement (F), and competition (G) assays expressed as CFU/mL. All assays at MOI 1:25, performed in triplicate with three independent biological replicates (n = 3). Data are mean ± SEM. Statistical analysis: one-way ANOVA with Dunnett’s multiple comparisons test. Significance: p < 0.05 (*), p < 0.01 (**), p < 0.001 (***), p < 0.0001 (****); ns, not significant.

### *L. johnsonii* attenuates MDR EAEC-induced gas production and nitric oxide synthesis but does not suppress EAEC growth through nutrient competition

To investigate the mechanistic basis of *L. johnsonii* antagonism beyond direct killing, gas production inhibition, macrophage nitric oxide modulation, and nutrient competition were assessed. In the gas production assay, MDR EAEC produced substantial fermentative gas evidenced by visible splitting of the LB soft agar layer and displacement of the overlying MRS+ agar, while *L. johnsonii* alone generated no detectable gas. Co-incubation with *L. johnsonii* markedly reduced agar splitting across three independent experiments, indicating suppression of EAEC fermentative metabolism, likely mediated by diffusible compounds secreted through the soft agar matrix (Figure 5A). To assess immunomodulatory activity, nitric oxide production was measured in MDR EAEC-infected RAW 264.7 macrophages using the Griess assay. MDR EAEC infection elevated nitrite concentrations approximately 39-fold above uninfected controls, confirming robust induction of macrophage-driven NO production. LPS stimulation produced comparable nitrite levels, validating the inflammatory model. Addition of *L. johnsonii* following EAEC infection significantly reduced nitrite levels by 67.7% (p < 0.05; Figure 5B), demonstrating attenuation of macrophage-derived inflammatory NO during MDR EAEC infection. To determine whether nutrient competition contributes to the observed anti-EAEC activity, *L. johnsonii* and MDR EAEC were co-cultured in both nutrient-rich McCoy’s 5A medium and nutrient-limited PBS. No significant differences in EAEC counts were observed under either condition (Figures 5C and 5D), indicating that nutrient competition does not account for the inhibitory activity of *L. johnsonii* observed in other assays and that the active mechanisms are primarily contact- or secretome-dependent rather than resource-driven.

**Figure 5.**
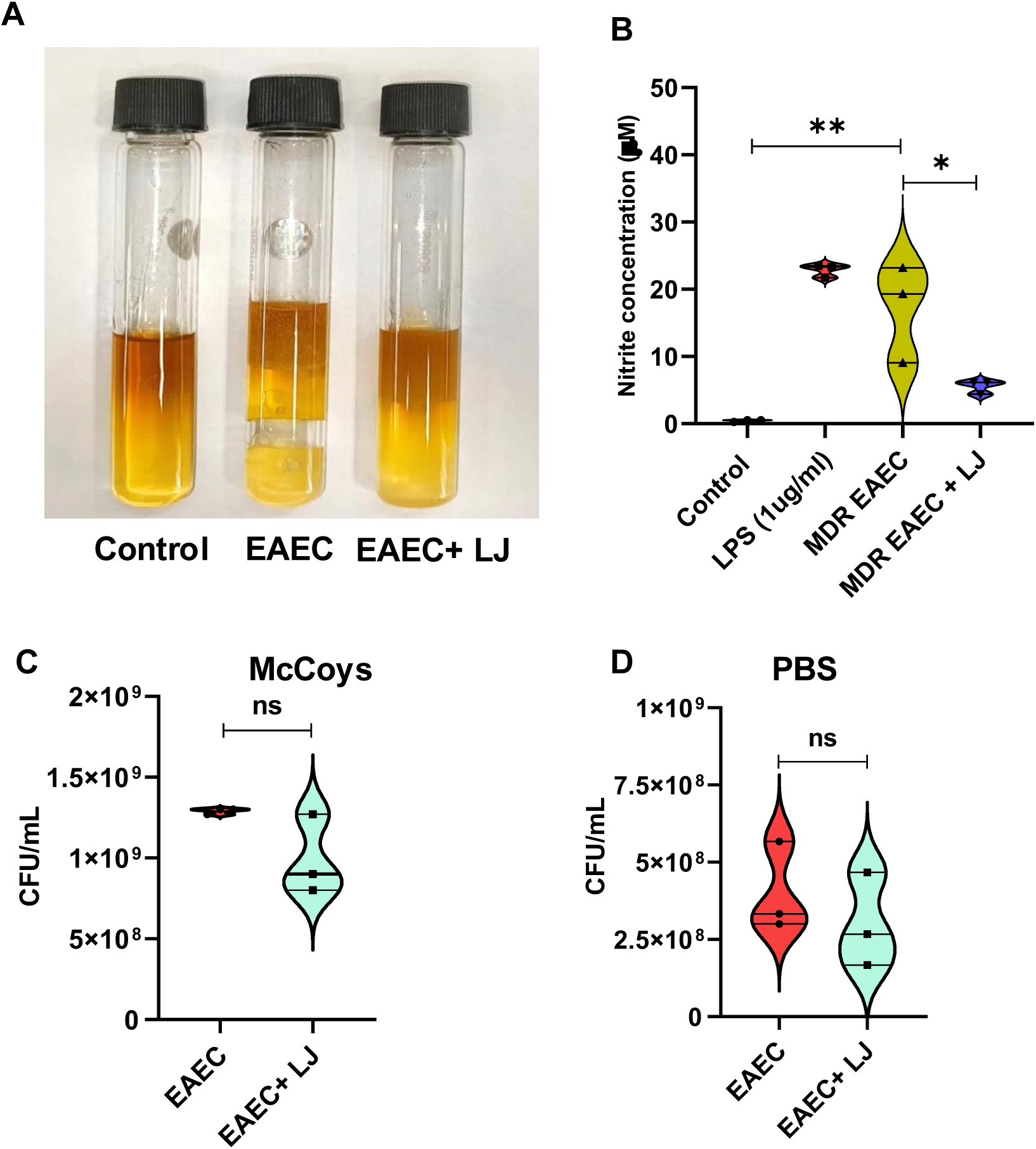
*L. johnsonii* attenuates MDR EAEC-induced gas production and macrophage nitric oxide synthesis but does not suppress EAEC growth through nutrient competition. (A) Gas production assay. MDR EAEC (1 × 10⁸ CFU) inoculated into 0.7% LB soft agar, overlaid with 0.7% MRS+ soft agar containing *L. johnsonii* (1 × 10⁸ CFU) or MRS+ alone. *L. johnsonii* in MRS+ overlaid with plain LB soft agar served as the probiotic gas control. Incubated at 37°C for 24 hours under anaerobic conditions. Gas production indicated by agar splitting or displacement. Image representative of three independent experiments. (B) Nitric oxide production by Griess reagent assay in RAW 264.7 macrophages infected with MDR EAEC at MOI 1:10 or stimulated with LPS (500 ng/mL) for 24 hours, followed by L. johnsonii at MOI 1:25 for 6 hours. Nitrite (µM) calculated from sodium nitrite standard curve. Uninfected untreated cells served as basal control. (C) Nutrient competition in McCoy’s 5A medium (nutrient-rich). MDR EAEC alone or co-cultured with *L. johnsonii* at 10⁷ CFU/mL each for 6 hours. MDR EAEC CFU/mL determined by plating on LB agar. (D) Nutrient competition in PBS (nutrient-limited) under identical conditions. All experiments performed in triplicate with three independent biological replicates (n = 3). Data are mean ± SEM. Statistical analysis: one-way ANOVA with Dunnett’s multiple comparisons test for panel B; unpaired Student’s t-test for panels C and D. Significance: p < 0.05 (*), p < 0.01 (**), p < 0.001 (***), p < 0.0001 (****); ns, not significant.

### *L. johnsonii* cell lysate and cell-free supernatant exhibit bactericidal activity against MDR EAEC, with sustained inhibition confined to FPLC fractions S5 and S6

To identify the molecular basis of *L. johnsonii* antimicrobial activity, cell lysate and cell-free supernatant (CFS) were tested against MDR EAEC at increasing concentrations, followed by FPLC fractionation of the CFS. Both preparations significantly inhibited MDR EAEC. The cell lysate produced significant reductions at 25 and 50 µg protein per 100 µL (46.5% and 43.3% respectively, p < 0.05), while 12.5 µg did not reach significance (Figure 6A). The CFS achieved significant inhibition only at 50 µg protein per 100 µL (55.1% reduction, p < 0.01), with lower concentrations showing trends that did not reach statistical significance (Figure 6B). The activity of the cell lysate at lower concentrations relative to the CFS suggests that cellular components contribute to antimicrobial efficacy alongside secreted factors. FPLC fractionation of the CFS on a Superdex 75 size-exclusion column yielded five fractions (S1, S2, S4, S5, and S6), all below 75 kDa (Figure 6C). At 1 hour, fractions S1, S2, S4, S5, and S6 all significantly inhibited MDR EAEC growth, with S6 producing the greatest reduction at this early time point (p < 0.001; Figure 6D). At 6 hours, however, only S5 and S6 retained significant inhibitory activity, reducing EAEC counts by 58.1% and 67.7% respectively (both p < 0.01), while S1, S2, and S4 lost activity (Figure 6E). The sustained bactericidal activity of S5 and S6 at 6 hours indicates the presence of stable bioactive molecules in these fractions, consistent with non-proteinaceous small-molecule effectors.

**Figure 6.**
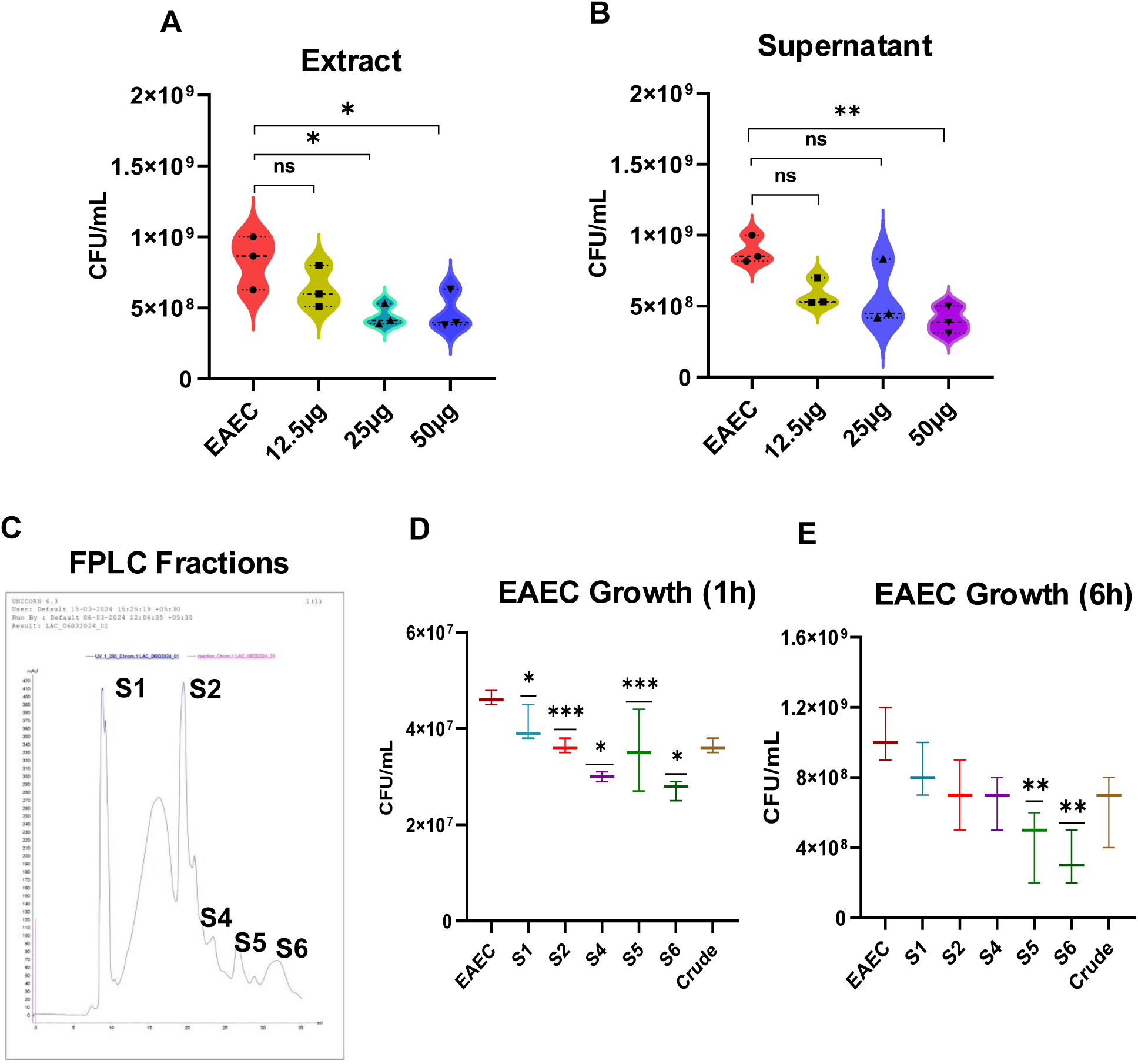
*L. johnsonii* cell lysate and cell-free supernatant exhibit bactericidal activity against MDR EAEC, with sustained inhibition confined to FPLC fractions S5 and S6. (A) Growth inhibition of MDR EAEC after 6 hours incubation with *L. johnsonii* cell lysate at 12.5, 25, and 50 µg protein per 100 µL. Untreated MDR EAEC served as the control. Results expressed as CFU/mL. (B) Growth inhibition of MDR EAEC after 6 hours incubation with *L. johnsonii* cell-free supernatant at 12.5, 25, and 50 µg protein per 100 µL. Results expressed as CFU/mL. (C) Representative FPLC chromatogram of *L. johnsonii* CFS fractionated on a Superdex 75 size-exclusion column (UV absorbance 280 nm). Five fractions (S1, S2, S4, S5, and S6) were collected below 75 kDa. Fraction S3 was not collected due to insufficient chromatographic resolution in that region. (D) Antimicrobial activity of FPLC fractions (S1, S2, S4, S5, S6) and crude CFS at 30 µg/mL against MDR EAEC after 1 hour incubation. Results expressed as CFU/mL. (E) Antimicrobial activity of the same fractions after 6 hours incubation. Results expressed as CFU/mL. All experiments performed in triplicate with three independent biological replicates (n = 3). Data are mean ± SEM. Statistical analysis: one-way ANOVA with Dunnett’s multiple comparisons test. Significance: p < 0.05 (*), p < 0.01 (**), p < 0.001 (***), p < 0.0001 (****); ns, not significant.

### Gram staining confirms progressive bactericidal activity of *L. johnsonii* CFS and differential killing efficacy of FPLC fractions S5 and S6 against MDR EAEC

To morphologically confirm and visually quantify the bactericidal activity of the CFS and active FPLC fractions, Gram staining was performed on MDR EAEC following treatment. CFS treatment produced a progressive, concentration-dependent reduction in visible EAEC cell counts of 32.7% at 25 µg/mL (p < 0.01) and 66.8% at 50 µg/mL (p < 0.0001) relative to untreated controls, with the remaining cells retaining intact rod morphology and uniform staining intensity (Figures 7A and 7B). These findings are consistent with the CFU data from Figure 6B and confirm that CFS reduces viable EAEC cell numbers in a dose-dependent manner.

**Figure 7.**
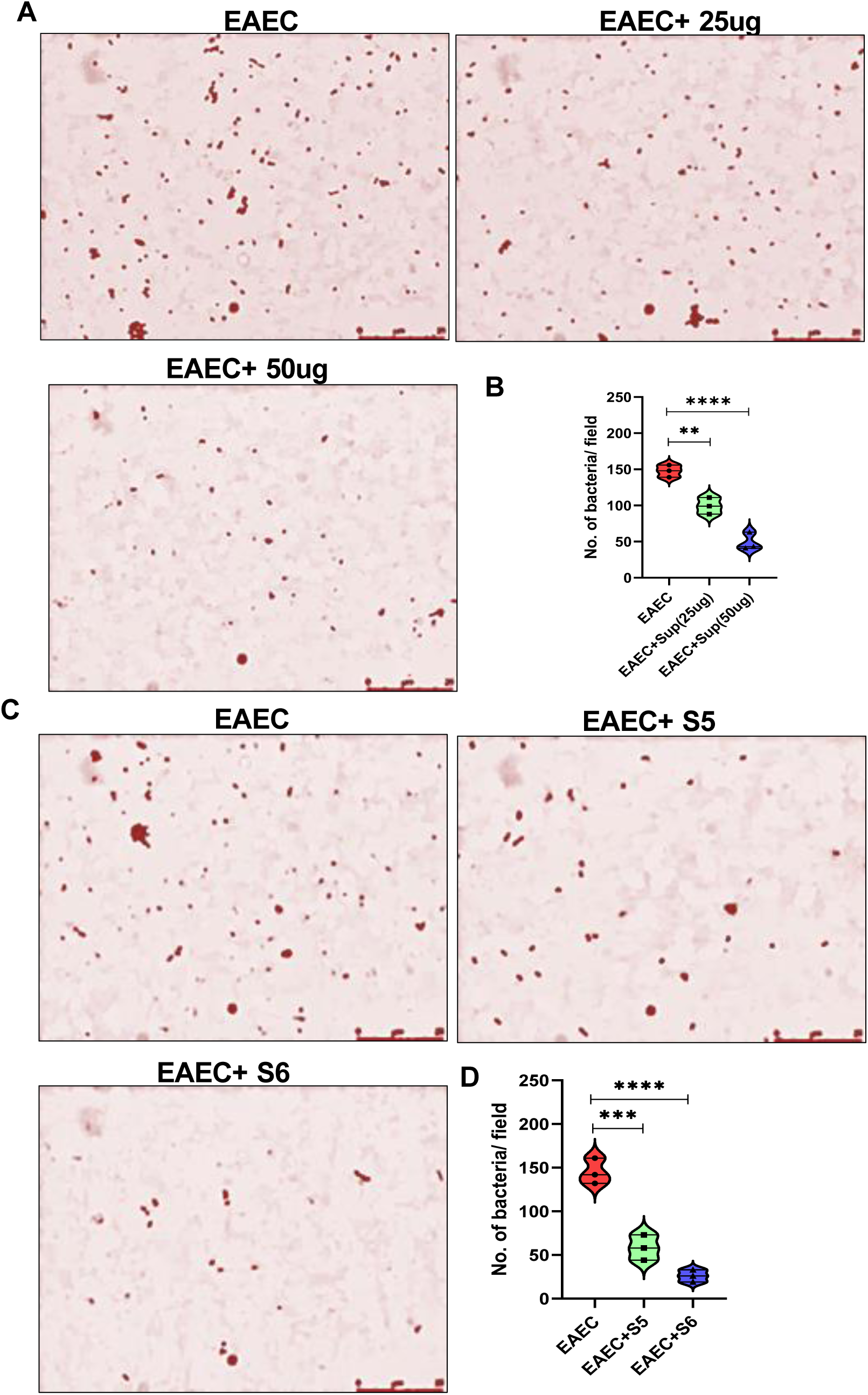
Gram staining confirms progressive bactericidal activity of *L. johnsonii* CFS and differential killing efficacy of FPLC fractions S5 and S6 against MDR EAEC. (A) Representative Gram-stained images of MDR EAEC following 6 hours treatment with *L. johnsonii* CFS at 25 µg/mL and 50 µg/mL. Untreated MDR EAEC served as the control. Oil immersion at 100× magnification. (B) Quantification of visible MDR EAEC cells per field following CFS treatment at 25 µg/mL and 50 µg/mL. CFS produced a concentration-dependent reduction of 32.7% at 25 µg/mL (p < 0.01) and 66.8% at 50 µg/mL (p < 0.0001) relative to untreated controls. Counts performed across a minimum of three fields per condition per experiment. Results expressed as number of bacteria per field. (C) Representative Gram-stained images of MDR EAEC following 6 hours treatment with fractions S5 and S6 at 30 µg/mL each. Untreated MDR EAEC served as the control. (D) Quantification of visible MDR EAEC cells per field following treatment with fractions S5 and S6 at 30 µg/mL each. S5 reduced visible cell counts by 59.8% (p < 0.001) and S6 by 82.1% (p < 0.0001) relative to untreated EAEC, indicating greater bactericidal potency of S6. Results expressed as number of bacteria per field. All images representative of three independent experiments. Data in panels B and D are mean ± SEM. Statistical analysis: one-way ANOVA with Dunnett’s multiple comparisons test. Significance: p < 0.01 (**), p < 0.001 (***), p < 0.0001 (****).

Treatment with fractions S5 and S6 produced quantitatively distinct outcomes. S5 reduced total visible EAEC cell counts by 59.8% relative to untreated EAEC (p < 0.001; Figures 7C and 7D), while S6 produced a more pronounced reduction of 82.1% (p < 0.0001), consistent with the greater bactericidal potency of S6 observed in the CFU-based assay at 6 hours (Figure 6E). The difference in killing magnitude between S5 and S6, sustained across both the CFU enumeration and Gram stain cell count readouts, indicates that these fractions represent distinct bioactive fractions with differing bactericidal efficacy. Characterization of the precise mechanism of action of each fraction will require future studies employing membrane permeability assays or electron microscopy.

## DISCUSSION

Enteroaggregative *Escherichia coli* is a major cause of persistent diarrhea in children in low- and middle-income countries, and the increasing prevalence of multidrug-resistant (MDR) strains has substantially limited treatment options^1, 2, 11^. The EAEC isolate used in this study exhibited resistance to multiple antibiotic classes, consistent with regional surveillance data, supporting the need for alternative therapeutic approaches^11^. In this context, the present study demonstrates that *L. johnsonii* exerts a broad anti-EAEC effect encompassing direct antimicrobial activity, biofilm disruption, competitive epithelial exclusion, suppression of gas production, attenuation of macrophage nitric oxide responses, and secretion of stable bactericidal components, collectively supporting its potential as a biotherapeutic candidate against MDR EAEC. Importantly, to our knowledge this is the first study to characterize fractionated secretome-derived bactericidal activity of *L. johnsonii* against MDR EAEC and to demonstrate its immunomodulatory effects in this pathotype context.

The probiotic characterization data indicate that *L. johnsonii* possesses the physiological attributes required for gastrointestinal survival and colonization. Its tolerance to acidic pH, bile salts, and digestive enzymes is consistent with previous CFU-based studies on this isolate^23^ and with prior characterization of *L. johnsonii* strains under simulated gastrointestinal conditions^29, 30^. Phenol tolerance serves as an established proxy for resistance to aromatic fermentation metabolites that accumulate in the colon under dysbiotic conditions^28^. High cell surface hydrophobicity and rapid autoaggregation reaching 80.4% by 4 hours indicate strong adhesion potential, critical for colonization and competitive exclusion^13^. These traits are particularly relevant in EAEC infection, where the pathogen relies on aggregation and biofilm formation to persist on the intestinal mucosa^3, 5^, making the competitive superiority of *L. johnsonii* in these traits mechanistically significant.

The antimicrobial assays demonstrate that *L. johnsonii* produces diffusible inhibitory factors capable of suppressing MDR EAEC. The agar overlay results, in which inhibition substantially exceeded that of gentamicin at 30 µg/mL, indicate strong secretome-mediated activity^31, 32^. Biofilm assays further revealed an 81.4% reduction in viable biofilm-associated EAEC, with live bacteria showing markedly greater efficacy than heat-killed cells, confirming that active metabolic processes are required for full anti-biofilm activity. This is consistent with previous reports linking *L. johnsonii* exopolysaccharides and secreted metabolites to biofilm interference^17, 18^. This viability dependence is consistent with reports by Miyazaki et al., who demonstrated that probiotic CFS activity against EAEC was pH-dependent and abolished upon neutralization, suggesting that metabolically active secretion of both organic acids and bacteriocin-like compounds contributes to the observed effect^33^. Notably, EAEC co-culture reduced *L. johnsonii* biofilm formation by 71.3%, suggesting reciprocal competitive interactions in the biofilm niche that may have practical implications for the timing of probiotic administration relative to infection establishment.

The protection assays revealed a time-dependent mechanism of epithelial colonization resistance. Pre-establishment of *L. johnsonii* reduced subsequent EAEC colonization by 59.6%, and post-infection addition displaced pre-adhered EAEC by 34.7%, while simultaneous co-inoculation produced no significant reduction. This pattern indicates that *L. johnsonii* requires prior epithelial occupancy to effectively exclude pathogens, likely through physical occlusion of adhesion sites and localized secretion of antimicrobial compounds at inhibitory concentrations, consistent with colonization resistance models described for other probiotic lactobacilli^34, 35^. This time-dependency is consistent with findings by Agbemavor et al., who demonstrated that pre-establishment of *L. plantarum* FS2 on epithelial surfaces more effectively inhibited diarrhoeagenic EAEC adhesion than competitive exclusion or displacement conditions^36^. These findings suggest that prophylactic administration of *L. johnsonii* may be more effective than therapeutic use after infection is established, and the enhanced probiotic proliferation observed under exclusion conditions further indicates that early colonization reinforces sustained pathogen suppression.

Beyond direct antimicrobial effects, *L. johnsonii* attenuated EAEC-associated physiological and inflammatory responses. Suppression of gas production suggests interference with EAEC fermentative metabolism, with implications for reducing gastrointestinal symptoms associated with infection^10^. The 67.7% reduction in nitric oxide production in infected macrophages is particularly relevant, as excessive NO contributes to mucosal oxidative damage and inflammation^37, 38^. This magnitude of reduction is comparable to that reported for conditioned media from *L. rhamnosus* GG in murine macrophages infected with *E. coli*^39^, and is likely mediated through modulation of NF-kB-driven inflammatory signaling^40^, positioning *L. johnsonii* as a dual-action immunobiotic. In contrast to its activity against EPEC and *C. rodentium*^23^, no nutrient competition-driven suppression of EAEC was observed under either nutrient-rich or nutrient-limited conditions, likely reflecting the metabolic versatility of EAEC, which encodes multiple biosynthetic and nutrient acquisition systems^8^. This pathotype-specific distinction indicates that anti-MDR EAEC activity is predominantly secretome-dependent.

FPLC fractionation identified fractions S5 and S6 as the principal bioactive components, both retaining sustained antimicrobial activity at 6 hours while S1, S2, and S4 lost activity, indicating the presence of stable low-molecular-weight compounds^31, 32^. Previous metabolomic characterization of this isolate identified fatty acyl derivatives, hydroxy acids, and alkaloids within these fractions^23^, consistent with the bactericidal activity confirmed here by Gram staining^41, 42^. S6 demonstrated greater bactericidal potency than S5 across both CFU enumeration and Gram stain cell count readouts, indicating that the *L. johnsonii* secretome contains multiple bioactive constituents with differing antimicrobial efficacies, consistent with observations by Mani-Lopez et al. that LAB cell-free supernatants harbor co-active antimicrobial components with distinct activity profiles against MDR pathogens^43^. These findings advance earlier work by Kumar et al.^44^and Miyazaki et al.^33^, who demonstrated probiotic activity against EAEC without identifying stable fractionable bioactives, by providing the first evidence of sustained bactericidal components below 75 kDa within a probiotic secretome that retain activity independently of pH-labile organic acids. The chemical identity and precise mechanism of action of both fractions remain unresolved and warrant future studies employing membrane permeability assays, electron microscopy, and MS/MS characterization.

This study has its own limitations. All experiments were conducted in vitro using a single EAEC isolate, and findings may not fully represent the diversity of EAEC strains^3, 4^. The use of a non-mucus-producing epithelial cell line may not capture the complexity of in vivo intestinal environments. The cell-free supernatant was not pH-neutralized, and while sustained activity in S5 and S6 argues against a purely pH-driven mechanism, the contribution of acidity at earlier time points cannot be fully excluded. Future studies should prioritize MS/MS characterization of S5, pH-controlled CFS experiments, evaluation of S5-S6 synergy, and in vivo validation to establish translational relevance and advance *L. johnsonii*-derived antimicrobials as viable alternatives in MDR EAEC-associated diarrheal disease.

## ACKNOWLEDGEMENTS

The authors gratefully acknowledge Dr. Sadhana Y. Veturi, Asian Institute of Gastroenterology (AIG Hospitals), Hyderabad, India, for providing the clinical MDR EAEC isolate used in this study under a formal Material Transfer Agreement. The authors also thank Dr. Prasada Rao H. B. D., National Institute of Animal Biotechnology, Hyderabad, India, for his valuable guidance and insights on fast protein liquid chromatography methodology.

## DATA AVAILABILITY STATEMENT

The raw and processed datasets underlying all figures and tables reported in this study are openly available in Zenodo at https://doi.org/10.5281/zenodo.19368379, including raw experimental data, analysis files, and presentation materials supporting the reported results. The 16S rRNA gene sequence of the *L. johnsonii* isolate used in this study was previously deposited in the NCBI GenBank nucleotide database under accession number PV739486 (https://www.ncbi.nlm.nih.gov/nuccore/PV739486) as part of a prior publication^23^ and is publicly accessible.

